# Whole-brain precision functional mapping of a proximal upper-extremity motor task

**DOI:** 10.64898/2026.09.22.752784

**Authors:** Neha A. Reddy, Michelle C. Medina, Ana Maria Acosta, Ahalya Mandana, Julius P. A. Dewald, Molly G. Bright

**Affiliations:** Department of Physical Therapy and Human Movement Sciences, Feinberg School of Medicine, Northwestern University, Chicago, IL, United States; Department of Biomedical Engineering, McCormick School of Engineering and Applied Sciences, Northwestern University, Evanston, IL, United States

**Keywords:** BOLD fMRI, motor task, precision functional mapping, shoulder abduction, whole brain

## Abstract

Upper-extremity motor control has been widely studied with hand-task functional magnetic resonance imaging (fMRI) in healthy and disease states. However, proximal-arm motor tasks, such as at the shoulder, are uncommon during fMRI due to practical limitations, though they are critical to a complete understanding of upper-extremity motor control. Previous fMRI studies of proximal upper-extremity control have generally used unconstrained movements or a single low force or torque condition, leaving unknown how distributed supraspinal brain activity scales with increasing proximal motor output. Here, we implemented a custom, MR-safe device to test supraspinal brain activity during three levels of isometric shoulder abduction in healthy individuals using a whole-brain fMRI acquisition. We conducted subject-specific analysis with repeated sampling of the controlled shoulder abduction task within each individual (i.e., “precision mapping”), as the exact localization of shoulder motor activity is relatively understudied. Our analysis showed spatial variability in subject-specific motor activation compared to group-level localization. We found that shoulder abduction had a significant effect on motor activity in the primary motor cortex, supplementary motor area, dorsal premotor area, thalamus, putamen, and cerebellum. Across repeated fMRI runs, amplitude of motor activity was consistent across all tested regions. Overall, these findings provide an individualized, whole-brain torque-response characterization of the proximal upper-extremity motor system and establish an approach for examining how distributed supraspinal motor networks are recruited as mechanical demand increases. These whole-brain, subject-specific methods can be a critical tool for probing motor dysfunction in clinical populations.

## Introduction

An understanding of the neural systems involved in upper-extremity motor control is critical to probing motor function and dysfunction. Several clinical conditions involve upper-extremity motor dysfunction, including stroke, Parkinson’s disease, and multiple sclerosis. A key tool in the study of whole-brain activity related to motor tasks is functional magnetic resonance imaging (fMRI). Importantly, fMRI allows for a non-invasive investigation of both surface and deep brain regions involved in motor control, including the cerebellum and basal ganglia. However, while hand-grasp tasks have been widely studied (Cramer, Weisskoff, et al., 2002; Hermsdörfer et al., 2003; Kuhtz-Buschbeck et al., 2008; Lindberg et al., 2012; Reddy, Zvolanek, et al., 2024; Ward et al., 2007), proximal-arm tasks, such as isometric shoulder abduction (SABD), are uncommon during fMRI. Importantly, SABD efforts may involve postural control and recruit different motor pathways compared to hand tasks (Davidson & Buford, 2004; Eilfort et al., 2025; Maslovat et al., 2023). In conditions such as stroke, SABD drives specific patterns of motor impairment (Lin et al., 2023; J. G. McPherson et al., 2018; L. M. McPherson & Dewald, 2022), requiring a detailed understanding of whole-brain SABD activity in healthy individuals to fully interpret.

The few fMRI studies conducted during proximal upper-limb tasks have been limited by their inability to control and modulate the motor tasks and have not examined whole-brain activity. For example, several groups instructed participants to perform elbow and shoulder movements during fMRI, though not at a specific angle or torque target (Broome et al., 2019; Gordon et al., 2023; Howard et al., 2019). Daly and colleagues (Daly et al., 2008) developed a device to allow participants to perform a stable reaching movement with the shoulder and elbow; however, this study also did not quantitatively control or measure the task. Krainak and colleagues (2007) successfully implemented an MR-safe device for performance of controlled proximal upper limb isometric tasks, but were limited to using a single low level of SABD torque and an event-related design to limit confounds caused by task-correlated head motion. None of these previous proximal arm movement studies investigated extra-cortical regions (Kulwattho et al., 2025), greatly limiting the full potential of fMRI to perform systems-level investigation of brain activity. Thus, it remains unknown how activity across the distributed human motor system scales with systematically increasing, individually normalized proximal-limb torque.

New fMRI methodological developments have been able to overcome the effects of head motion in motor-task fMRI and implement whole-brain analyses. Our previous work implemented multi-echo independent component analysis (ME-ICA) to provide more reliable estimates of motor activity even in datasets with high amounts of task-correlated head motion (Reddy et al., 2025; Reddy, Zvolanek, et al., 2024). Other task-fMRI studies have also supported the use of ME-ICA to improve sensitivity and mitigate effects of task-related head motion (Gonzalez-Castillo et al., 2016; Lombardo et al., 2016; Steel et al., 2022). The capabilities of ME-ICA expand the possibilities of the shoulder tasks implemented by Krainak and colleagues and allow for a study of controlled proximal motor tasks across varied torque levels. We also successfully implemented an fMRI protocol to study whole-brain activity during sensory and auditory tasks, demonstrating that whole-brain analyses are possible and feasible with fMRI data (Medina, Reddy, Sitek, et al., 2026; Reddy, Clements, et al., 2024).

An additional challenge in proximal upper-limb fMRI is that standard group-level analyses may not be sufficient to appropriately characterize motor activity. Compared to hand-grasp activity, which has been well localized to the hand-knob region of the motor cortex (Yousry et al., 1997), shoulder activity is relatively understudied and may not produce reliable results at the group level. In recent years, several resting-state and task-fMRI studies have underscored the utility of a precision functional mapping approach, characterizing subject-specific functional regions and connectomes, to overcome the obscuring effects of group averaging (Gordon et al., 2017, 2023; Marek & Greene, 2021). ME-ICA has also been shown to improve reliability of precision functional mapping results, particularly in subcortical regions (Lynch et al., 2020, 2021; Medina, Reddy, Sitek, et al., 2026). Subject-specific ME-ICA analyses have the potential to better localize proximal motor activity and probe individual variability.

Here, we use an MRI-compatible six-degree-of-freedom load cell and multi-echo fMRI techniques to measure motor activation in healthy individuals who performed a controlled SABD task at three torque levels. We take a subject-specific precision functional mapping approach to understand shoulder motor activation and collect densely sampled shoulder motor-task data in each subject. We aim to characterize subject-specific SABD activity across varied torque levels in motor regions throughout the whole brain. We hypothesized that activity within cortical, putaminal, thalamic, and cerebellar motor regions would increase with normalized SABD torque, providing a systems-level characterization of the neural recruitment required for progressively greater proximal motor output. As a secondary analysis, we compare localization of SABD motor activity to motor activity during a well-characterized hand grasp task to better understand variability between subjects.

## Methods

### Data collection

#### Participants

This study was approved by the Northwestern University Institutional Review Board, and all participants provided written, informed consent. Seven right-handed individuals with no known history of neurological or vascular disorders participated (4M, 23 ± 1y). Prior to MRI scanning, each participant underwent collection of maximum SABD torque in an isometric setup. MRI scanning was completed over two days, during which participants performed an isometric unilateral SABD task with their non-dominant (i.e., left) arm. During MRI session one, participants completed the SABD task at Low (14 ± 2% maximum), Medium (35 ± 5% maximum), and High (56 ± 9% maximum) torque. During MRI session two, participants completed an additional SABD task at Medium (35 ± 5% maximum) torque. One participant (sub-05) was not able to participate in session two.

#### Collection of maximum torques

Participants’ left-arm maximum voluntary torques (MVTs) were collected on a separate day from the MRI sessions to prevent fatigue from influencing MRI results. In order to collect the most accurate maximum values, participants were secured in a chair with torso straps (Biodex Inc.). A previous study demonstrated that SABD torques collected in the seated position are not significantly different from those collected in the supine position required by the MRI environment (Krainak et al., 2011).

Participants’ maximum SABD torques were collected using a six degree-of-freedom (DOF) load cell (JR3 Inc.). Participants were instructed to perform isolated SABD tasks without involvement of the trunk. A forearm orthosis secured the participant’s arm firmly to the load cell, which was placed in the plane of the forearm and located approximately midway between the wrist and the elbow (See Figure 1). The highest value of five maximum torque trials was recorded as the participant’s maximum torque for use in the MRI sessions; if the participant’s fifth trial was the highest, additional trials were collected until a clear maximum torque was identified.

**Figure 1.**
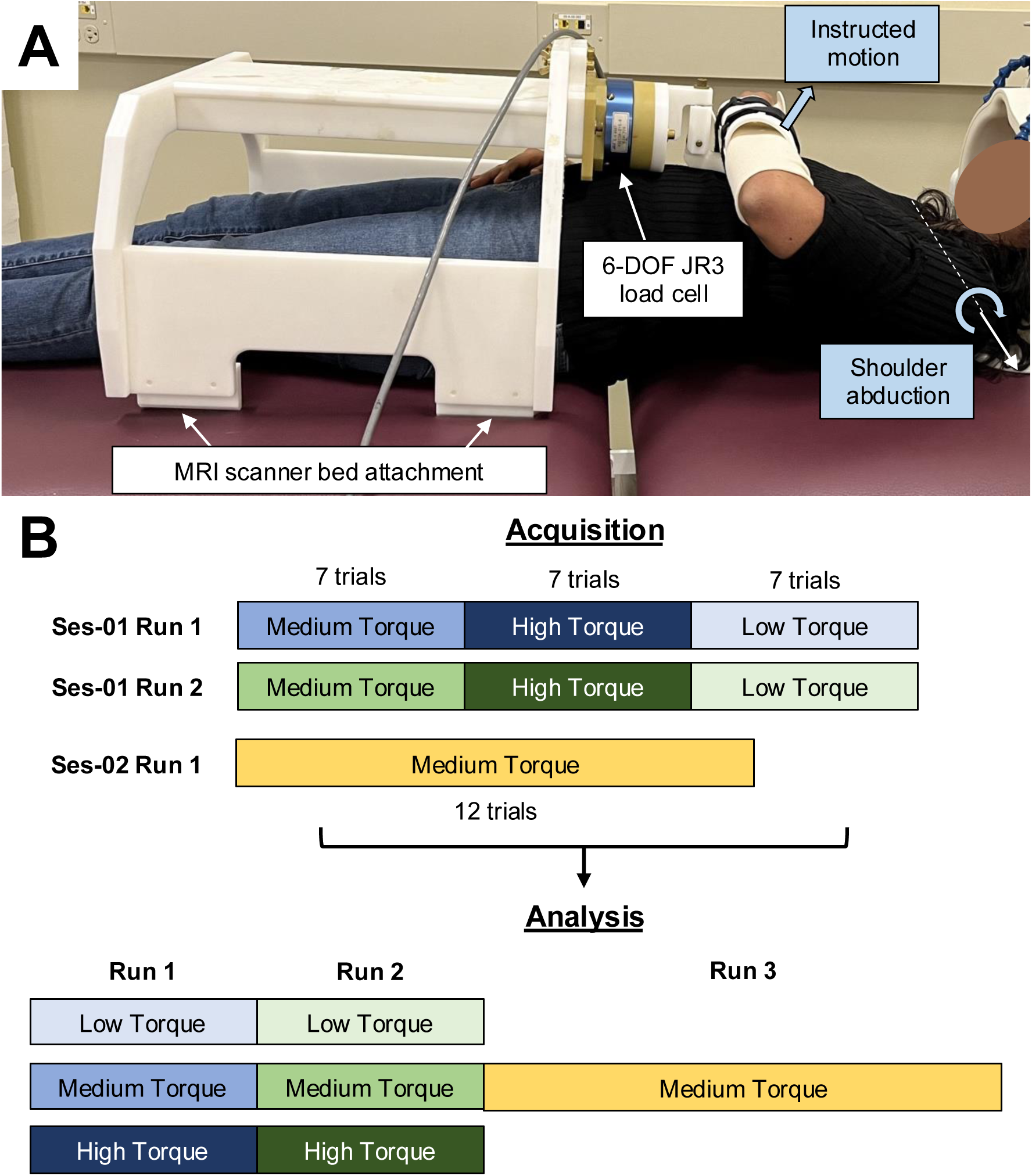
(A) Shoulder abduction measurement device. (Author N.A.R. pictured) (B) Acquisition and analysis of shoulder abduction data across two MRI sessions.

#### Scan protocol

Participants were scanned on a Siemens 3T Prisma MRI system with a 32-channel head coil. During session one, a structural T1-weighted multi-echo MPRAGE image was collected using parameters adapted from Tisdall and colleagues (Tisdall et al., 2016): TR = 2.17 s, TEs = 1.69/3.55/5.41 ms, TI = 1.16 s, FA = 7°, FOV = 256 x 256 mm^2^, and voxel size = 1 x 1 x 1 mm^3^. The three echo images were combined using root-mean-square.

During both sessions, functional scans were collected using a multiband multi-echo gradient-echo echo planar imaging sequence provided by the Center for Magnetic Resonance Research (CMRR, Minnesota): TR = 2.2 s, TEs = 13.4/39.5/65.6 ms, FA = 90°, MB factor = 2, GRAPPA = 2, voxel size = 1.731 x 1.731 x 4 mm^3^, 44 slices, phase encoding direction A >> P, field of view 180 mm, matrix size 104 x 104 (Moeller et al., 2010; Setsompop et al., 2012). Axial slices were aligned perpendicular to the base of the fourth ventricle to maximize both in-plane resolution in the brainstem and whole-brain coverage (Reddy, Clements, et al., 2024). A reverse-phase encode P >> A scan was acquired before each functional scan for use in distortion correction. CO_2_ was collected throughout functional scans via nasal cannula, measured at 1000 Hz sampling rate.

#### SABD tasks

During MRI session one, participants used a device with an MR-safe six-DOF load cell (JR3 Inc.) (Krainak et al., 2007) (Figure 1) to perform an isometric left-arm SABD task targeting Low, Medium, and High SABD torque: 10-s “lift”, 15-s “relax”. Participants performed the motor task during two functional scans (runs 1 and 2) with seven consecutive trials per torque level; each scan consisted of a total of 21 motor task trials during 250 functional volumes (9 minutes and 10 seconds). Order of torque levels was randomized across participants and kept consistent between functional scans one and two within each participant.

During MRI session two, participants used the same device to perform the isometric left-arm SABD task at Medium torque, only. During the functional scan (run 3), participants performed 12 motor task trials over 201 functional volumes (7 minutes 22 seconds); each trial was 10-s “lift”, 15-s “relax”. After every 4 trials, there was an additional 1-minute rest period.

During both sessions, participants viewed instructions and real-time torque feedback on a screen. Text on the screen instructed the start and end of the task periods. During the task periods, a box with height of target torque ± 5% MVT indicated the target torque level. A moving bar indicated the participants’ real-time torque level and turned green when the torque was within the target range. Force and torque signals in six DOF were collected from the load cell throughout the scans, measured at 1000 Hz sampling rate. A passive auditory stimulus was presented simultaneously during motor-task periods, as part of a separate study (Medina, Reddy, Sitek, et al., 2026).

#### Hand-grasp task

To compare SABD motor activity to a more established distal motor task, we collected hand-grasp data in two participants for use in a secondary analysis. During MRI session two, two participants (sub-01 and sub-04) performed an isometric left hand-grasp task during an additional fMRI scan. The hand-grasp device consisted of a one-DOF load cell (Interface, Inc.) fixed between two halves of a Delrin rod for participant comfort. Before the start of the scan, the SABD measurement device was removed from the scanner. The hand-grasp device was positioned in the participant’s left hand, with the arm placed in a resting position by the participant’s side. During the functional scan, participants performed 12 motor trials over 201 functional volumes; each trial was 10-s “squeeze”, 15-s “relax”. After every 4 trials, there was an additional 1-minute rest period. The target grasp force was 25% of the participant’s maximum voluntary contraction (MVC), collected before the scan using a Jamar Hand Dynamometer. The highest value of five maximum force trials was recorded as the participant’s maximum force for use in the MRI sessions; if the participant’s fifth trial was the highest, additional trials were collected until a clear maximum force was identified. Visual task instructions were identical to those of the SABD tasks. Hand-grasp force from the load cell was collected throughout the scan, measured at 100 Hz sampling rate.

### Data analysis

#### Structural MRI pre-processing

T1-weighted images for each subject were processed with FSL’s (Jenkinson et al., 2012) fsl_anat, which performs bias field correction and brain extraction.

#### Functional MRI pre-processing

FSL (Jenkinson et al., 2012) and AFNI (Cox J.S., 1996) tools were used for fMRI preprocessing. The first 10 volumes of each echo time series were removed to allow for steady-state magnetization to be attained; then distortion correction was performed (topup, FSL). For MRI session one scans, each scan was divided into three segments by torque level, which were analyzed independently. Head-motion realignment was estimated for the first echo data, with reference to the Single Band reference image taken at the start of the scan (3dVolreg, AFNI), and then applied to all echo time series (3dAllineate, AFNI). All images were brain extracted (bet, FSL). Tedana (Ahmed et al., 2024; Dupre et al., 2021) was used to calculate a T2*-weighted combination of the three echo datasets, producing the optimally combined (ME-OC) fMRI dataset. Multi-echo independent component analysis (ME-ICA) was performed on the ME-OC fMRI data using tedana. The resulting components were automatically classified with tedana’s external regressors decision tree (Ahmed et al., 2024); components that were correlated (p < 0.05, R^2^ > 0.5) with input nuisance regressors (CSF grey matter timeseries and motion parameters) were rejected. ME-OC timeseries at each voxel were converted to signal percentage change for further analysis, then data were smoothed at 3mm FWHM (3dmerge, AFNI).

#### Motor-task and end-tidal CO_2_ regressors

##### Motor task

The six degree-of-freedom (DOF) signals from the load cell were transformed to the shoulder joint using measured participant arm lengths / angles. The real-time SABD torque was convolved with a canonical double-gamma hemodynamic response function, rescaled from 0 to 1, downsampled to the resolution of the functional MRI images (TR = 2.2s) using the volume triggers from the MRI scan, then demeaned. The temporal derivatives of these task regressors were used to account for small subject- and region-specific differences in hemodynamic response. A version of the task regressors was also made that was normalized by the participant’s MVT (regressor units were converted to % MVT). For each subject-level model described below, two task regressors were used: (1) the real-time-torque-derived regressor and (2) its derivative.

##### End-tidal CO_2_

End-tidal peaks were detected in the CO_2_ data using an automatic peak finder in MATLAB, then manually inspected. The end-tidal peaks were interpolated to form an end-tidal CO_2_ trace that was convolved with a canonical hemodynamic response function, rescaled to the range of the unconvolved timeseries, downsampled to the resolution of the functional MRI images (TR = 2.2s) using the volume triggers from the MRI scan, then demeaned.

#### Subject-level models to localize motor activity

All SABD scans were modeled together to optimize localization of shoulder activity: 2 runs of Low torque, 2 runs of Medium torque, and 2 runs of High torque from session one, and 1 (longer) run of Medium torque from session two. The session two scan was transformed into the functional space of session one before further analysis, using a concatenated transform of session two functional to anatomical space, then anatomical to session one functional space (epi_reg, FLIRT, FSL). As sub-05 did not participate in session two, only the 6 SABD runs from session one were incorporated for this subject’s analysis. For each subject, the ME-OC signal from each voxel was processed using a general linear model (GLM) that incorporated all functional scan runs and included six motion parameters and their derivatives from volume realignment, up to fourth-order Legendre polynomials, the % MVT motor-task regressor and its derivative, the end-tidal CO_2_ regressor, and the rejected ME-ICA components (AFNI, 3dREMLfit). The rejected ME-ICA components were conservatively orthogonalized to the two motor-task regressors, end-tidal CO_2_ trace, motion parameters, Legendre polynomials, and accepted ME-ICA components, as done previously (Moia et al., 2021; Reddy et al., 2025). An F-test for the combination of the two motor-task regressors was run using the -gltsym option. This F-test allowed for consideration of slight variability in the timing of the motor response across subjects and motor regions.

#### Group-level analysis

Group-level analysis was performed to provide a comparison to the subject-specific results. For each subject, the individual beta coefficient maps of the motor-task regressor and its derivative from subject-level localization analysis were both converted to MNI space by applying a concatenated transform of functional to anatomical space (FLIRT and FNIRT, FSL) and anatomical to functional space (epi_reg, FLIRT, FSL). Group-level analysis was performed with AFNI’s 3dMVM (Chen et al., 2014), using a contrast for task + derivative > 0 (Harms & Melcher, 2003; Henson & Friston, 2007). Group-level maps were thresholded at p < 0.005 and clustered at alpha < 0.05, using a right-sided t-test (3dFWHMx, 3dClustSim, 3dClusterize, AFNI).

#### Creation of subject-specific ROIs

Subject-specific ROIs were determined in the right upper-extremity primary motor cortex (M1), right supplementary motor area (SMA), right dorsal premotor (PMd), right thalamus, right putamen, and left cerebellum. For the right upper-extremity M1 ROI, a mask of the right upper-extremity M1 from the Brainnetome atlas, unthresholded, (Fan et al., 2016) was dilated by two voxels to create a conservative mask, then transformed to each subject’s functional space using a concatenated spatial transformation of the MNI-space mask to the subject’s T1-w structural image (FLIRT and FNIRT, FSL) and then from this image to the subject’s functional images (epi_reg with field map unwarping, FSL). The top 10% of F-statistics within the ‘robust range’ (2 to 98%) in the upper-extremity M1 mask were considered to be the subject-specific ROIs (fslmaths, FSL). This method allowed for ROI creation independent of each map’s magnitude of F-statistics, as done previously (Medina, Reddy, Bright, et al., 2026; Medina, Reddy, Sitek, et al., 2026).

ROIs were similarly created within undilated masks of the right SMA from the Human Motor Area Template atlas (Mayka et al., 2006), unthresholded; the right PMd from the Brainnetome atlas, unthresholded; the right thalamus from the Harvard-Oxford subcortical atlas (Frazier et al., 2005), thresholded at 50%; the right putamen from the Harvard-Oxford subcortical atlas, thresholded at 50%; and the left cerebellum from the MNI structural atlas (Collins et al., 1995; Mazziotta et al., 2001), thresholded at 50% and four midsagittal slices removed to mitigate noise.

For use as a negative control, a Heschl’s gyrus auditory ROI was similarly created within a mask of the left and right Brodmann’s areas 41 and 42 from the Brainnetome atlas, unthresholded.

#### Motor activation across SABD torque levels

To assess the magnitude of activation within each ROI across SABD torque levels, subject-level models were run separately for scans from each torque level using the previously described real-time-torque-derived task regressor rescaled from 0 to 1 and its derivative (*Motor-task and end-tidal CO_2_ regressors*). For each subject and torque level, the ME-OC signal from each voxel was processed using a general linear model that incorporated two functional scan runs (for Low and High torque) or three functional scan runs (for Medium torque) and included six motion parameters and their derivatives from volume realignment, up to fourth-order Legendre polynomials, the motor-task regressor and its derivative, the end-tidal CO_2_ regressor, and the rejected, orthogonalized ME-ICA components (AFNI, 3dREMLfit).

In order to calculate a beta coefficient that considered motor activation related to both the motor-task regressor and its derivative, we first reconstructed a motor activation timeseries for each voxel as *β_stimulus_* ∗ *Timeseries_stimulus_* + *β_derivative_* ∗ *Timeseries_derivative_*. As the task regressors were scaled 0 to 1, we calculated the range of the motor activation timeseries as the reconstructed voxel-wise combined motor beta coefficient.

For each subject and torque level, the median combined motor beta coefficient was calculated within the subject-specific ROIs described above. The effect of SABD (% MVT) on the measured activation was modeled as BetaCoefficient ∼ SABD + (1|Subject) using the *lme4* package in R (Bates et al., 2015), and significance of SABD was assessed using Satterthwaite’s method for approximating denominator degrees of freedom, as implemented in *lmerTest* in R (Kuznetsova et al., 2017). Pairwise comparisons were also performed between Low, Medium, and High SABD torque levels. SABD activity was modeled as BetaCoefficient ∼ TorqueLevel + (1|Subject), and estimated marginal means were calculated using *emmeans* in R (Lenth & Piaskowski, 2026). Bonferroni correction was used to control for multiple comparisons.

To compare the use of subject-specific ROIs to those from group-level results, we also created a set of ROIs from group-level analysis. The intersections of the group-level significant clusters with the right M1, right SMA, left cerebellum, and bilateral Heschl’s gyrus atlas regions were found and transformed to each subject’s functional space. The median combined motor beta coefficient was calculated within each of these ROIs for each subject and torque level. The effect of SABD (% MVT) on the measured activation and pairwise comparisons between Low, Medium, and High SABD torque levels were calculated as above.

#### Motor activation across repeated runs of SABD

To assess differences in activation between repeated runs of the same SABD task, subject-level models were individually calculated for the Medium torque scans from each run (run1, run2, and run3). As sub-05 did not participate in session two, only run1 and run2 were calculated for this subject’s analysis. The task regressors used were the previously described real-time-torque-derived task regressor rescaled from 0 to 1 and its derivative (*Motor-task and end-tidal CO_2_ regressors*). For each subject and run, the ME-OC signal from each voxel was processed using a general linear model that incorporated one functional run and included six motion parameters and their derivatives from volume realignment, up to fourth-order Legendre polynomials, the motor-task regressor and its derivative, the end-tidal CO_2_ regressor, and the rejected, orthogonalized ME-ICA components (AFNI, 3dREMLfit). A combined motor activation beta coefficient was calculated as above.

For each subject and run, the median combined motor beta coefficient was calculated within the subject-specific ROIs described above. The effect of Run on the measured activation was modeled as BetaCoefficient ∼ Run + (1|Subject) using the *lme4* package in R (Bates et al., 2015), and significance of Run was assessed using Satterthwaite’s method for approximating denominator degrees of freedom, as implemented in *lmerTest* in R (Kuznetsova et al., 2017). Using the same model, pairwise comparisons were also performed between run1, run2, and run3. Estimated marginal means were calculated using *emmeans* in R (Lenth & Piaskowski, 2026). Bonferroni correction was used to control for multiple comparisons.

#### Hand-grasp task

For the two participants who performed the hand-grasp task, subject-level GLM analysis and ROI creation were conducted in an identical manner to the subject-level SABD analysis described in *Subject-level models to localize motor activity*, incorporating only one hand-grasp fMRI run per subject.

## Results

All SABD maximum torque values, structural MRI scans, and functional MRI scans were successfully collected as described above. As sub-05 did not complete MRI session two, one fewer functional MRI scan (Medium torque SABD task) was incorporated into subject-level analysis. SABD torques during each task are shown in Supplemental Table 1. Real-time SABD torque measurements are shown in Supplemental Figure 1. Participants were able to match the three target SABD torque levels well during task periods. However, during some rest periods, participants did not completely relax their arm, particularly sub-01 and sub-02.

### Subject-specific ROIs versus group-level clusters

Subject-specific ROIs for regions of greatest shoulder activation were created in the right upper-extremity M1, right SMA, right PMd, right thalamus, right putamen, and left cerebellum. The overlap of the seven subject-specific ROIs in each of these regions is shown in Figure 2. Significant clusters of activation from group-level analysis are overlaid in green outline. Some group-level clusters are related to activation from the concurrent auditory stimuli. Overall, the significant group-level clusters did not align well with regions of greatest subject-level shoulder activation overlap, and the exact spread of each ROI differed across regions. In the right M1, the ROIs converged on a specific region that was adjacent to a group-level cluster. In the left cerebellum, the ROIs converged on specific regions that corresponded with a group-level cluster, with the largest overlap occurring in lobules I-IV, V, and VIIIa. The spread of the ROIs for each subject varied considerably more in the cerebellum compared to in the M1. In the right SMA and right putamen, subjects converged on a specific region, though there were no voxels where all seven subjects overlapped. Group-level analysis did not yield any clusters in these regions. In the right thalamus and PMd, the subject-specific ROIs similarly demonstrated considerable variability and no group-level clusters were detected. An example of the subject-level variability in the right thalamus and PMd is shown in Figure 3. For each region, subject-specific ROIs from three subjects are shown, with little overlap in foci of motor activation between subjects.

**Figure 2.**
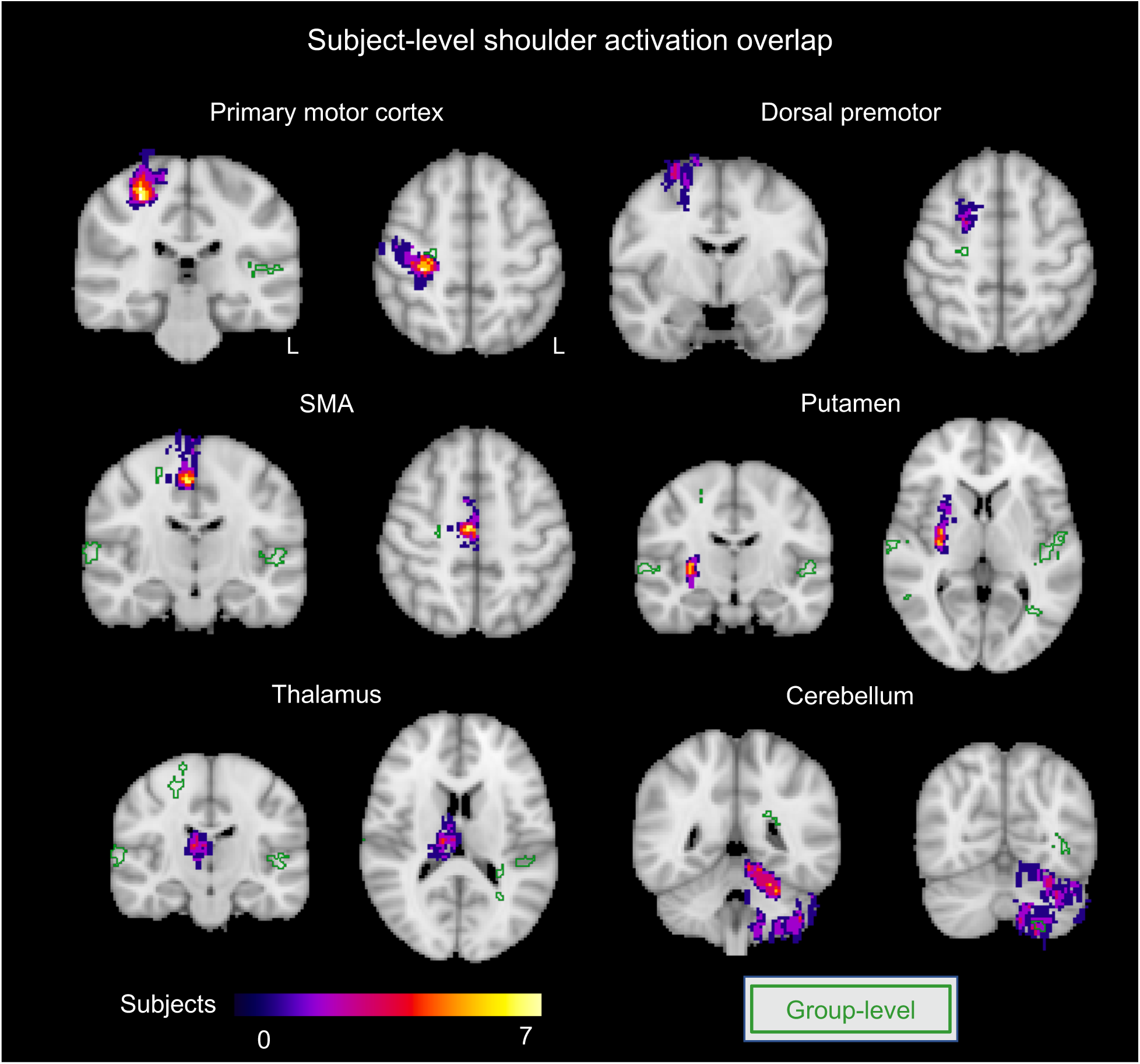
Subject-level activation related to shoulder motor activity. Seven subjects were included. The top 10% of F-statistics in each ROI were transformed to MNI space for each subject, then results from all subjects were overlaid. For comparison, the group-level results for shoulder motor activity are shown in green outline.

**Figure 3.**
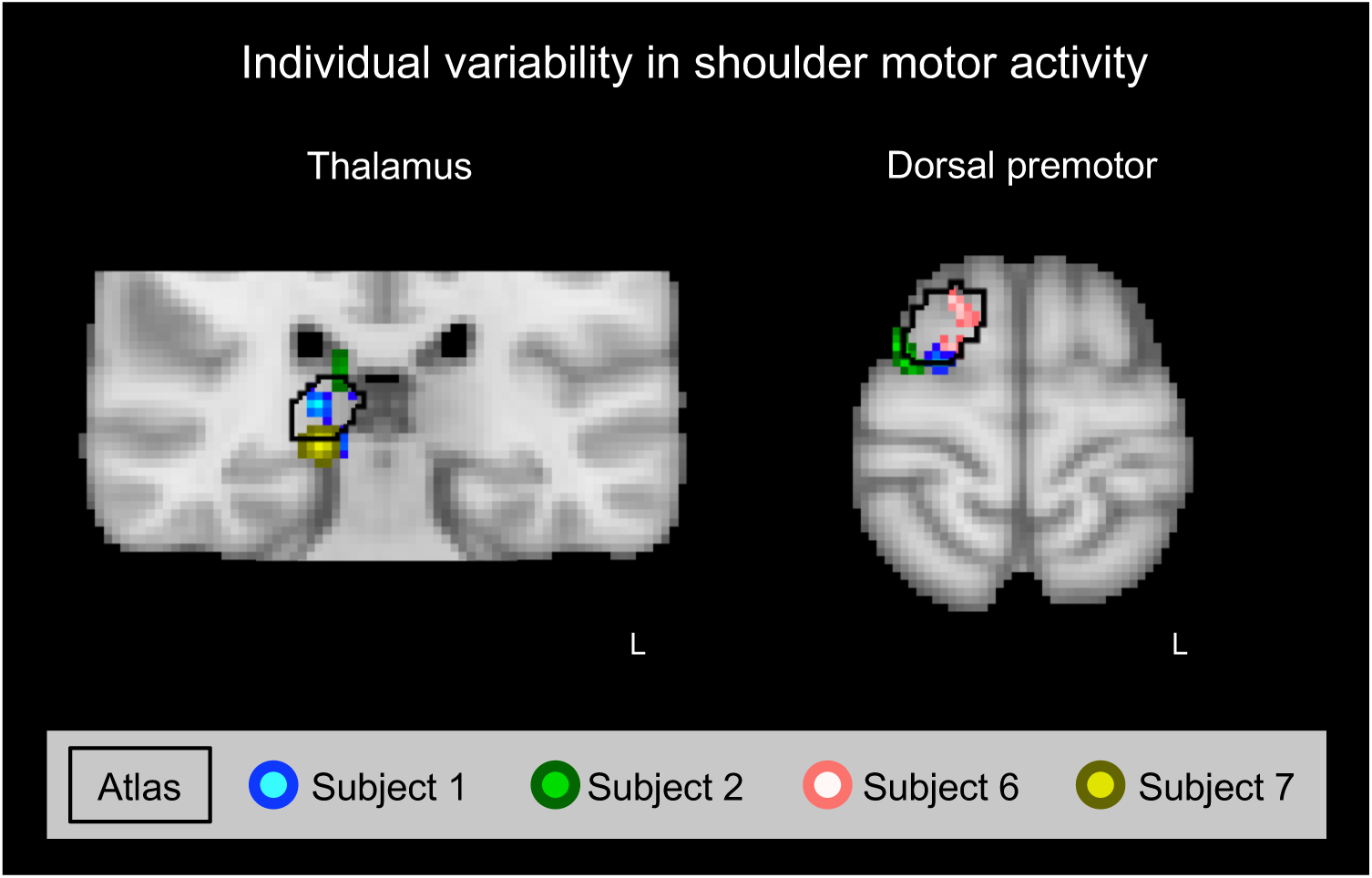
Individual variability in shoulder motor activity in the right thalamus and dorsal premotor (PMd) regions. For the thalamus and PMd, motor activity from three subjects in each region are shown, as well as the atlas regions used to create the subject-level ROIs. There is little overlap between the main foci of activation for each subject shown.

### Motor activation across three SABD torque levels

The amplitude of motor activation for each torque level was determined as the median combined motor beta coefficient within each subject-specific motor ROI and the Heschl’s gyrus, which served as a negative control. We expected the amplitude of motor activation to increase with higher torque levels in motor regions. The median beta coefficient values (as percent signal change) for each subject, ROI, and SABD value (as % MVT) are shown in Figure 4. The same results grouped by categorical torque level (Low, Medium, High) are shown in Supplemental Figure 2. We found a significant effect of SABD (% MVT) on percent signal change in all motor regions considered: right M1 (p < 0.0005), right SMA (p < 0.0001), right PMd (p < 0.0005), left cerebellum (p < 0.001), right putamen (p < 0.001), right thalamus (p < 0.001). Regarding Bonferroni-corrected pairwise comparisons in the categorical data, the M1, SMA, and cerebellum had a significant increase (p < 0.05) from Low to Medium and from Low to High torque. The PMd, thalamus, and putamen only showed a significant increase (p < 0.05) between Low and High torque levels. In the Heschl’s gyrus, as expected, we found no effect of torque (% MVT) on percent signal change (p = 0.51), with no significant difference in signal change between each categorical torque level.

**Figure 4.**
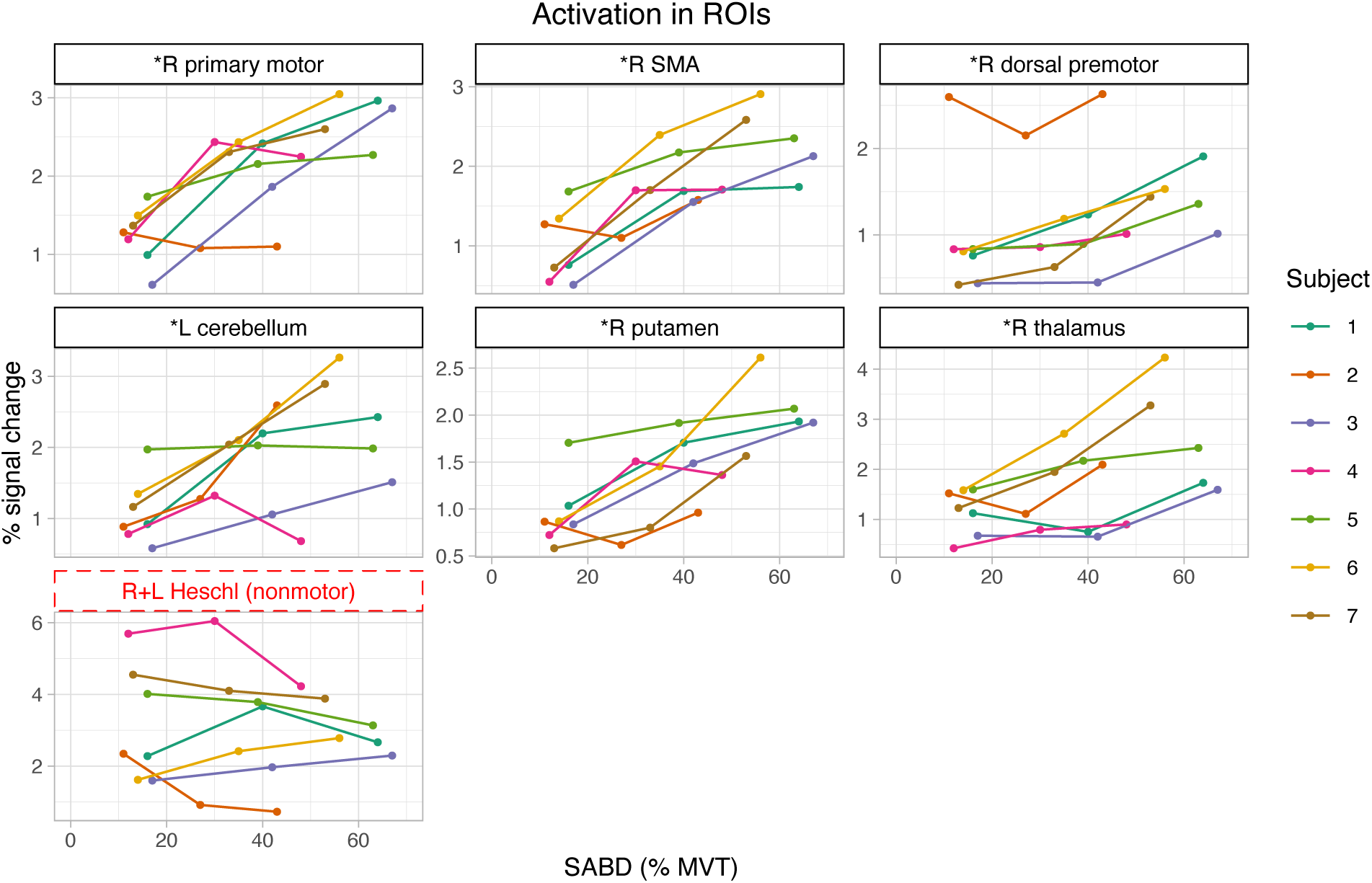
Activation across torque levels in motor and non-motor ROIs. The top 10% of F-statistics in each motor region was calculated to create subject-specific ROIs. All motor ROIs had a significant effect of SABD (% MVT) on signal change, indicated by an asterisk next to ROI name (BetaCoefficient ∼ SABD + (1|Subject)). There was no significant effect of SABD on signal change in the Heschl’s gyrus, related to auditory activation, which was expected to have consistent activation across all motor tasks.

The amplitude of motor activation for each torque level was also calculated within ROIs created from group-level results (Supplemental Figure 3), to probe agreement with the subject-specific ROIs. (Note, only three motor regions and Heschl’s gyrus were identified as having significant group-level activation clusters.) We found a significant effect of SABD (% MVT) on percent signal change in the right M1 (p < 0.005) and left cerebellum (p < 0.05). We did not find a significant effect of percent signal change in the right SMA (p = 0.52) or the Heschl’s gyrus (p = 0.76). The M1 showed a significant increase (p < 0.05) from Low to Medium and from Medium to High torque. There was no significant difference in signal change between each torque level in the SMA, cerebellum, or Heschl’s gyrus.

### Motor activation across three Medium torque runs

We compared the amplitude of motor activation across the three Medium torque runs for each subject to determine consistency of calculated percent signal change. Sub-05 only had two Medium torque runs, as they did not complete MRI session two. Run1 and run2 were obtained during a single MRI session. Run3 was collected during a second MRI session on a different day. We did not see a significant effect of Run on percent signal change, or a difference in signal change between torque levels, in any ROI (Figure 5). This finding indicates that the calculated percent signal change was fairly stable across runs, even on different days.

**Figure 5.**
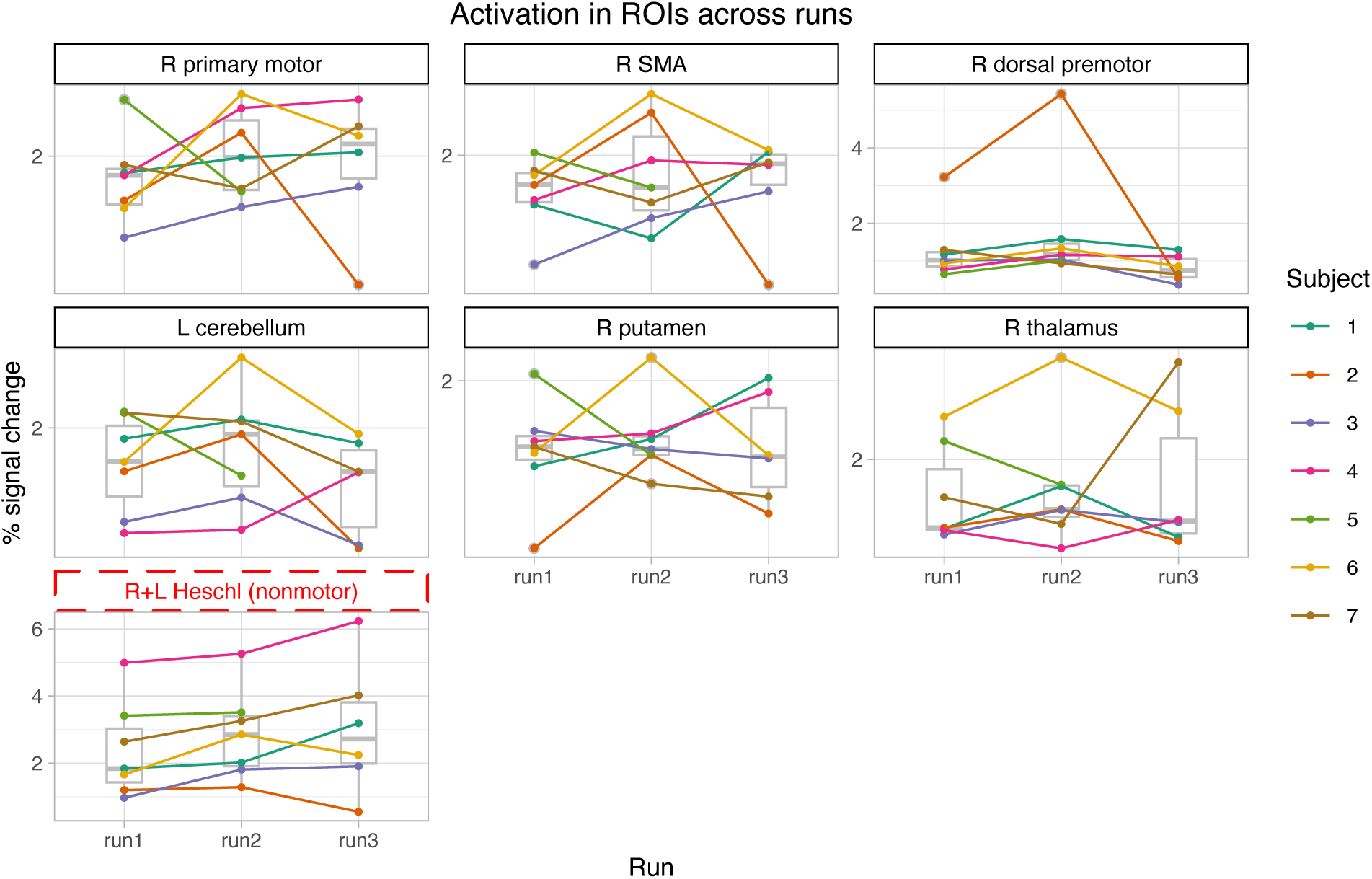
Consistency of motor activation across runs of the Medium torque task for each subject. Run1 and run2 were obtained during a single MRI session. Run3 was collected during a second MRI session on a different day. There were no ROIs with a significant effect of Run on signal change (BetaCoefficient ∼ Run + (1|Subject)).

### Hand-grasp motor activity

We collected hand-grasp data from two subjects to compare an established distal motor task to subject-specific SABD data (Figure 6). We found that sub-04 demonstrated a linear motor organization in M1, with SABD activity more medial to hand-grasp activity; sub-01 potentially demonstrated a concentric organization, with two foci of SABD activity surrounding hand-grasp activity. In the SMA, PMd, and cerebellum, our results again show variability between subjects.

**Figure 6.**
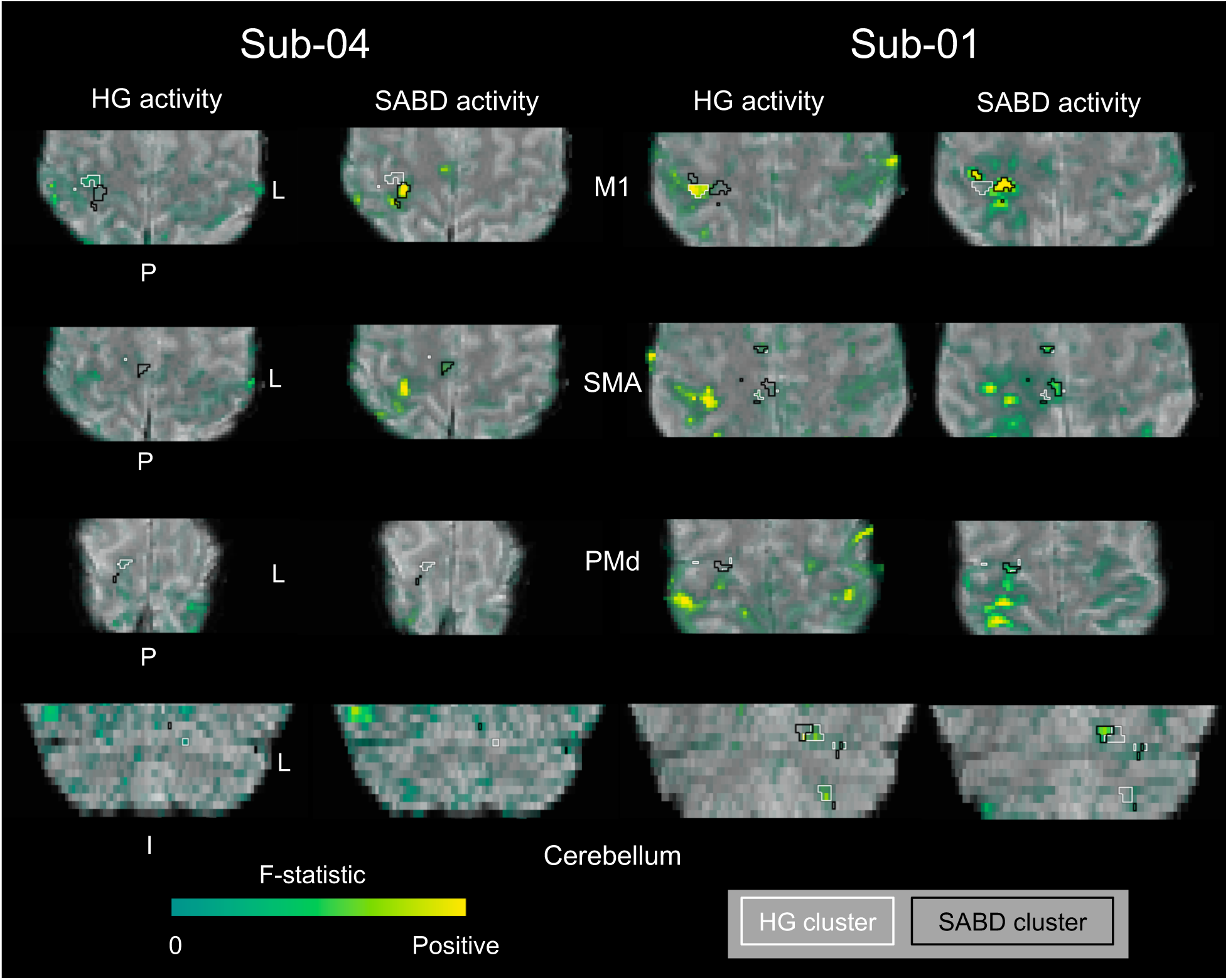
Hand-grasp (HG) and shoulder abduction (SABD) cortical activity for two representative subjects, with contralateral M1, SMA, and PMd and ipsilateral cerebellum. F-statistic maps are shown, with opacity modulated by the F-statistic magnitude. The F-statistic range for each map was selected separately to best highlight contrast across each map. The top 10% of F-statistics within a mask of each motor region is outlined in white for HG and black for SABD.

## Discussion

We used a controlled SABD motor task at three torque levels to identify subject-specific motor activity across expected cortical and subcortical brain regions. We identified subject-specific regions of motor activity in the contralateral M1, SMA, PMd, thalamus, putamen, and ipsilateral cerebellum. There was a significant effect of SABD on amplitude of motor activity in all of these motor regions. We also found that the amplitude of activity did not vary significantly across repeated fMRI runs in all regions tested.

### Subject-specific SABD motor activity was identified in expected cortical and subcortical brain regions

In this study, for the first time, we demonstrated the successful implementation of a controlled, isometric SABD task at varied torque levels in the MRI environment. Overall, we found that participants matched the three target SABD torque levels well (Supplemental Figure 1), consistently maintaining a torque of ± 5% target. Sub-01 and sub-02 did not always return to a neutral torque during rest periods. We implemented real-time torque tracings into our GLM analysis to account for this and other discrepancies in task timing compared to measured SABD torques.

We expected to detect motor activity in several cortical and subcortical motor areas: contralateral M1, SMA, PMd, thalamus, putamen, and ipsilateral cerebellum. The roles of these regions in healthy motor control are well established. The contralateral M1 is the key cortical region driving the corticospinal tract, the major descending motor pathway for voluntary movement. The SMA and PMd also contribute to the corticospinal tract and contain projections to M1. Compared to M1, SMA and PMd are known to play a greater role in proximal upper-extremity and trunk motor control, compared to distal upper-extremity motor control, and contribute to the corticoreticulospinal tract (Li et al., 2019). The SMA and PMd are also thought to play a role in motor planning and learning (Nachev et al., 2008; Vanderah & Gould, 2025). The thalamus, putamen, and cerebellum are involved in feedback loops that modulate movement (Vanderah & Gould, 2025).

We determined subject-specific ROIs in each of the above motor regions to better characterize activity related to SABD. Overall, proximal arm motor tasks have not been as well characterized as hand motor tasks. Though hand grasp is localized to the hand knob area of the primary motor cortex (Yousry et al., 1997), shoulder motor tasks do not have a specific known anatomical correlate in cortical or other regions. Critically, group results may mask individual motor activity due to variability across subjects, which is the motivation for many precision mapping studies (Gordon et al., 2017, 2023; Lynch et al., 2020, 2021; Marek & Greene, 2021; Medina, Reddy, Sitek, et al., 2026). Here, we implemented a precision mapping approach to fully understand individual proximal motor activity and compared our results to those of a traditional group-level analysis. Near the M1 and cerebellum clusters identified in group-level analysis, we found largely overlapping but variable subject-specific activity (Figure 2). Group-level analysis failed to fully capture the extent of individual subject motor activity in these regions, as its main strength is capturing common regions of activity across subjects. In the SMA, group-level analysis captured only one voxel of significant motor activity; and in the PMd, putamen, and thalamus, it did not capture any significant activity (Figure 3). When we examined subject-specific activation in these areas, we found significant variability in location of activation. Therefore, insufficient overlap of subject-level activation likely contributed to lack of group-level clusters in the SMA, PMd, putamen, and thalamus. While lower strength of motor signal change in a region can also contribute to failure of group-level analysis to identify clusters, our results demonstrated similar amplitude of activation in the SMA, PMd, putamen, and thalamus compared to the M1 and cerebellum (Figure 4). It is important to note that slight differences in location between subjects may result from transformation of results from functional to standard space, though we used a well-established method (FLIRT, FNIRT, FSL). These differences make co-localization of small foci of activation imperfect (Medina, Reddy, Sitek, et al., 2026). Our findings highlight the importance of a precision mapping approach in understanding whole-brain activation during proximal upper-extremity motor tasks.

### SABD motor activity is associated with torque level in cortical and subcortical brain regions

We aimed to understand how motor activity in each region changed with increasing SABD. To avoid the confounds of non-overlapping activation across subjects, and the non-significant activation at the group-level in select motor regions, we calculated activation amplitude using the subject-specific ROIs. We found a significant effect of SABD on motor activity in all tested motor regions (Figure 4). Our negative control, the non-motor auditory cortex, was not correlated with torque level, suggesting that our findings are not simply a result of increased task-correlated artifact, such as head motion, during production of greater torques. Our results are aligned with previous studies of hand grasp showing that cortex, thalamus, putamen, and cerebellum activity are positively correlated with force (Cramer, Weisskoff, et al., 2002; Dai et al., 2001; Keisker et al., 2009; Kuhtz-Buschbeck et al., 2008; Spraker et al., 2007; Wasson et al., 2010). We also performed a similar analysis using regions derived from group-level analysis. We found that the group-level results identified a significant effect of SABD on motor activity in the M1 and cerebellum, but failed to do the same in the SMA, a region with more variable subject-level activity (Supplemental Figure 3). Even when the subject-level data from each torque level was input separately to increase the apparent sample size (21 versus 7 inputs to the model), group results demonstrated fewer significant differences between torque levels compared our subject-specific approach (Supplemental Figure 4, Supplemental Figure 5). This result again underscores the benefits of a subject-specific approach when characterizing motor activity.

### Motor activity across scan sessions was consistent in all motor regions

We found that the amplitude of motor activity was consistent across fMRI runs within the same scan session and across different days in the M1, SMA, PMd, thalamus, putamen, cerebellum, and Heschl’s gyrus (Figure 5). This finding is encouraging for studies of subject-specific motor activity and/or clinical populations, in which larger quantities of data may be necessary to achieve study goals. Collecting data across different motor runs and scan sessions may also help to reduce the effects of fatigue on data (Liu et al., 2003; van Duinen et al., 2007), which participants with motor impairment may be particularly prone to experiencing (Kuppuswamy et al., 2015). Liu and colleagues (Liu et al., 2003) found that fatigue was associated with increased activity in cortical and cerebellar regions, while van Duinen and colleagues (van Duinen et al., 2007) saw decreased activation in SMA. In another study of the thalamus and basal ganglia, these regions were also associated with decreased activation during periods of fatigue (Hou et al., 2016). Our findings demonstrate that scans can be collected during different fMRI sessions if desired to minimize fatigue, without compromising consistency of shoulder motor activation results.

### Location of subject-specific motor activity may differ across individuals

We found variability in location of subject-specific motor activity across participants, particularly in the PMd and thalamus (Figure 3). Our results suggest that predefined atlases and group-level findings may not be sufficient to accurately localize proximal upper-extremity motor activity in healthy individuals. In addition, studies that focus on group-level findings might not detect significant activity in all motor regions, even when motor activity exists on the subject level (Figure 2). Higher sample sizes can help improve group-level detection of shoulder motor activity (Supplemental Figure 4). Deepening our knowledge of motor variability in healthy individuals is critical to interpretation of findings in diseased states; results may be due to normal variation in physiology or due to pathology.

As a secondary analysis, we sought to compare our subject-specific localization of upper-extremity motor activity to previous findings by including a left hand grasp task for two subjects. Within-subject comparisons also undergo fewer transformations during analysis. Previous fMRI studies of M1 have reported results that are aligned with a linear organization (Alkadhi et al., 2002; Plow et al., 2010) and others aligned with a concentric organization of proximal limbs surrounding distal limbs (Gordon et al., 2023; Meier et al., 2008). In favor of a linear organization, Alkhadi and colleagues (Alkadhi et al., 2002) found that participants demonstrated upper-extremity M1 somatotopy in a gradient from hand to wrist to elbow. In favor of a concentric organization, Meier and colleagues (Meier et al., 2008) found that forearm and wrist activity surrounded hand activity in M1, but did not find the same result for elbow movements. Gordon and colleagues (Gordon et al., 2023) tested several tasks in a precision functional mapping experiment with two individuals and found more compelling evidence that elbow and shoulder activity may also surround hand activity.

Here, we implemented a controlled motor task and whole-brain analysis to evaluate motor somatotopy in M1 and other motor regions. Results from two subjects showed variable motor organization in M1, SMA, PMd, and cerebellum (Figure 6). Gordon and colleagues (Gordon et al., 2023) also demonstrate variability between their two precision-mapped subjects who performed motor tasks. In the context of previous findings, our results and those of Gordon and colleagues suggest that there may be significant subject-level variability in macroscopic upper-extremity M1 organization. Some subjects may demonstrate activity more aligned with the “linear” organization model and some with the “concentric” model, and this may manifest in other motor regions outside of M1. Future studies of motor activity in a larger dataset are important to further explore concentric patterns of activation, critically, with a subject-specific analysis approach.

### Limitations

Though we were able to test an isometric SABD task with our device, isolation of specific muscles was not ensured or measured. We expected our SABD task to primarily require activation of the shoulder abductor muscles, mainly the anterior deltoid. It is possible that participants concurrently activated muscles at the elbow, wrist, hand, and trunk during the task. In fact, as elbow motor activity is also concurrently measured with our 6-DOF load cell, we observed that many participants produced a simultaneous elbow flexion or extension during the shoulder task (Supplemental Figure 6). The higher amplitude torque tasks may have also led to recruitment of a broader range of muscles. Therefore, our observation of greater amplitude brain activity during higher shoulder torques may have been a result of both recruitment of greater motor units within the primary SABD muscles and greater recruitment of motor units within other muscles. Future studies that wish to disentangle these effects and precisely measure muscle activity during task performance can consider concurrent electromyography (EMG) during fMRI scanning. However, this technique is uniquely challenging and is associated with separate experimental limitations (Ganesh et al., 2007).

The amount of fMRI data collected in our study is also a potential limitation. While our analysis approach allowed us to identify subject-specific ROIs for all participants, sub-05 did not complete session two and had less fMRI data used for analysis. Overall, sub-05’s results did not show a clear increase in motor activity with greater torque levels, compared to other participants (Figure 4), even in cortical regions. This suggests that sub-05 may not have had enough data to produce reliable motor ROIs. Sub-05 was also the participant with the lowest SABD MVT, greater than 1 standard deviation lower than the mean (Supplemental Table 1). Our motor task was normalized to the participants’ MVTs to capture Low, Medium, and High effort, rather than absolute torque. However, notably, sub-05 and sub-02, the participants with the two lowest SABD MVT values, were the only two participants to not demonstrate a clear increase in motor activity with greater torque levels in M1, which has been well-established to correlate positively with increased effort (Cramer, Weisskoff, et al., 2002). The interaction of absolute SABD torques with amplitude of brain activity, both mechanistically and as it relates to our specific experimental design, is a topic for further exploration. While determining the amount of data required to produce reliable subject-level motor ROIs was not a goal of this study, we note that all participants other than sub-05 had just under 26 minutes of fMRI data, with increasing signal change with torque level seen across multiple ROIs. 26 minutes is comparable or greater than the amount of data collected per motor task in other precision functional mapping studies (Gordon et al., 2017, 2023).

Another consideration is the method of ROI creation we implemented in this study. We determined subject-level ROIs as the top 10% of F-statistics within the ‘robust range’ (2 to 98%) of each motor region. Previous work in task fMRI analysis showed that this method yielded more consistent activation clusters in the thalamus and cerebellum compared to voxel-wise and cluster-based approaches (Medina, Reddy, Bright, et al., 2026). However, our ROI-creation method may also be more prone to the effects of noise and artifact in voxels distant from the center of activation. While our ROI-creation approach was effective in identifying the primary regions of activation for each subject, collection of more data may enable detection of ROIs with cluster-based approaches that better mitigate effects of noise. However, cluster-based approaches have limited spatial specificity, which may obscure detection of smaller ROIs and nuclei (Woo et al., 2014).

We calculated an F-statistic to test for the combined effects of a motor-task regressor and its derivative. Our analysis method allowed us to account for slight variabilities in the motor response timing across subjects and motor regions. For example, we observed that the motor response in the PMd was slightly earlier than the response in the M1 (Supplemental Figure 7). One potential reason for this timing variability is differences in the hemodynamic response across brain regions (Lewis et al., 2018). Future work may wish to further investigate these timing differences or consider implementing more advanced analysis techniques.

### Considerations for future work

Our work demonstrates the feasibility of subject-specific identification of motor activity during SABD of up to ∼60% maximum voluntary torque. (Dewald et al., 1995; J. G. McPherson et al., 2018; L. M. McPherson & Dewald, 2022). Notably, this study is the first to show the association between torque and brain activity during proximal upper-extremity motor tasks, which are more difficult to study with fMRI due to practical limitations. The ability to study graded-torque proximal motor tasks is particularly important in certain clinical populations, including chronic hemiparetic stroke. While hand grasp tasks have previously been used to study post-stroke motor control (Cramer, Mark, et al., 2002; Ward et al., 2006, 2007), these studies necessarily exclude participants who have motor impairments that affect the hand. Distal function, including finger and hand movements, is often affected in individuals with severe stroke-related motor impairment (Lin et al., 2023). Therefore, these individuals may be excluded from standard distal motor task fMRI studies due to an inability to perform the task adequately. A SABD task can be performed by a wider range of participants and enables the study of whole-brain activation in post-stroke individuals with moderate to severe motor impairment.

In addition, SABD has been shown to be coupled with elbow flexion in post-stroke participants, driving the ‘flexion synergy’ across the affected limb (Dewald et al., 1995; J. G. McPherson et al., 2018; L. M. McPherson & Dewald, 2022). The ability to study proximal upper-extremity motor activity using fMRI broadens the scope of what we can learn about post-stroke motor reorganization, particularly in subcortical areas that cannot be studied with electroencephalography (EEG). In addition, our multi-echo whole-brain protocol enables simultaneous detection of activity across cortical, subcortical, and cerebellar regions, while mitigating task-correlated head movements, making systems-level analysis feasible in clinical cohorts. Studying post-stroke whole-brain activity during SABD is critical for a deeper understanding of the contributions of specific brain regions and descending motor pathways to the flexion synergy.

A subject-specific approach to determining ROIs is also particularly important in the study of stroke and other clinical impairments, as individuals may demonstrate variable brain activity depending on the location of their infarct or disease process. As individuals with post-stroke motor impairment exhibit greater head motion during motor tasks (Reddy, Zvolanek, et al., 2024; Seto et al., 2001), it is important to note that the maximum torque level that can be tested and effectively denoised is likely lower than that in healthy individuals. However, multi-echo analyses have been shown to improve denoising in similar populations and motor tasks (Reddy et al., 2025; Reddy, Zvolanek, et al., 2024).

Future studies may also wish to further explore subject-specific differences in localization of upper-extremity motor tasks. Our methods enable a controlled, isometric method of studying shoulder motor tasks, with the potential to test elbow tasks as well. Combined with a well-established isometric hand grasp task, analyzing fMRI data during hand, elbow, and shoulder motor tasks would help to investigate potential somatotopic variability between individuals.

## Conclusions

In this study, we implemented an isometric left-arm shoulder abduction task at three different torque levels with task-fMRI. We localized subject-specific motor activity in key regions throughout the brain: primary motor cortex (M1), supplementary motor area (SMA), dorsal premotor area (PMd), thalamus, putamen, and cerebellum. We found a significant effect of torque level on amplitude of motor activity in all motor regions. Amplitude of motor activity was also stable across different fMRI runs in all regions tested here. We observed that the localization and distribution of shoulder and hand motor activity differed between two individuals. These findings in healthy individuals are important for future studies of clinical populations, which may benefit from subject-specific and whole-brain approaches to studying motor control of the entire upper limb.

## Author Contributions

Neha A. Reddy: Conceptualization, Methodology, Software, Formal analysis, Investigation, Data curation, Writing - original draft, Writing - review & editing, Visualization, Project administration. Michelle C. Medina: Conceptualization, Methodology, Software, Formal analysis, Investigation, Writing - review & editing. Ana Maria Acosta: Methodology, Software, Writing - review & editing. Ahalya Mandana: Methodology, Software, Writing - review & editing. Julius P. A. Dewald: Conceptualization, Methodology, Writing - review & editing, Supervision, Funding acquisition. Molly G. Bright: Conceptualization, Methodology, Writing - review & editing, Supervision, Project administration, Funding acquisition.

## Acknowledgements

This work was supported by the National Institute of Child Health and Human Development at the National Institutes of Health (R03HD113915), the National Institute of Biomedical Imaging and Bioengineering at the National Institutes of Health (T32EB025766), and the American Heart Association (25PRE1356822 to M.C.M.). The content is solely the responsibility of the authors and does not necessarily represent the official views of the National Institutes of Health. This work was supported by the Center for Translational Imaging at Northwestern University and through the computational resources and staff contributions provided for the Quest high performance computing facility at Northwestern University, which is jointly supported by the Office of the Provost, the Office for Research, and Northwestern University Information Technology.

