## Supplemental Materials for "Whole-brain precision functional mapping of a proximal upper-extremity motor task"

Supplemental Table 1. Maximum voluntary torque (MVT) during left arm shoulder abduction and targeted percentage of MVT during Low, Medium, and High torque tasks for each subject.

| Subject | MVT (Nm) | Low (% MVT) | Medium (% MVT) | High (% MVT) |
|---|---|---|---|---|
| **1** | 30.0 | 16 | 40 | 64 |
| **2** | 9.1 | 11 | 27 | 43 |
| **3** | 19.2 | 17 | 42 | 67 |
| **4** | 10.8 | 12 | 30 | 48 |
| **5** | 6.4 | 16 | 39 | 63 |
| **6** | 32.9 | 14 | 35 | 56 |
| **7** | 32.8 | 13 | 33 | 53 |
| **Mean** | 20 ± 12 | 14 ± 2 | 35 ± 6 | 56 ± 9 |

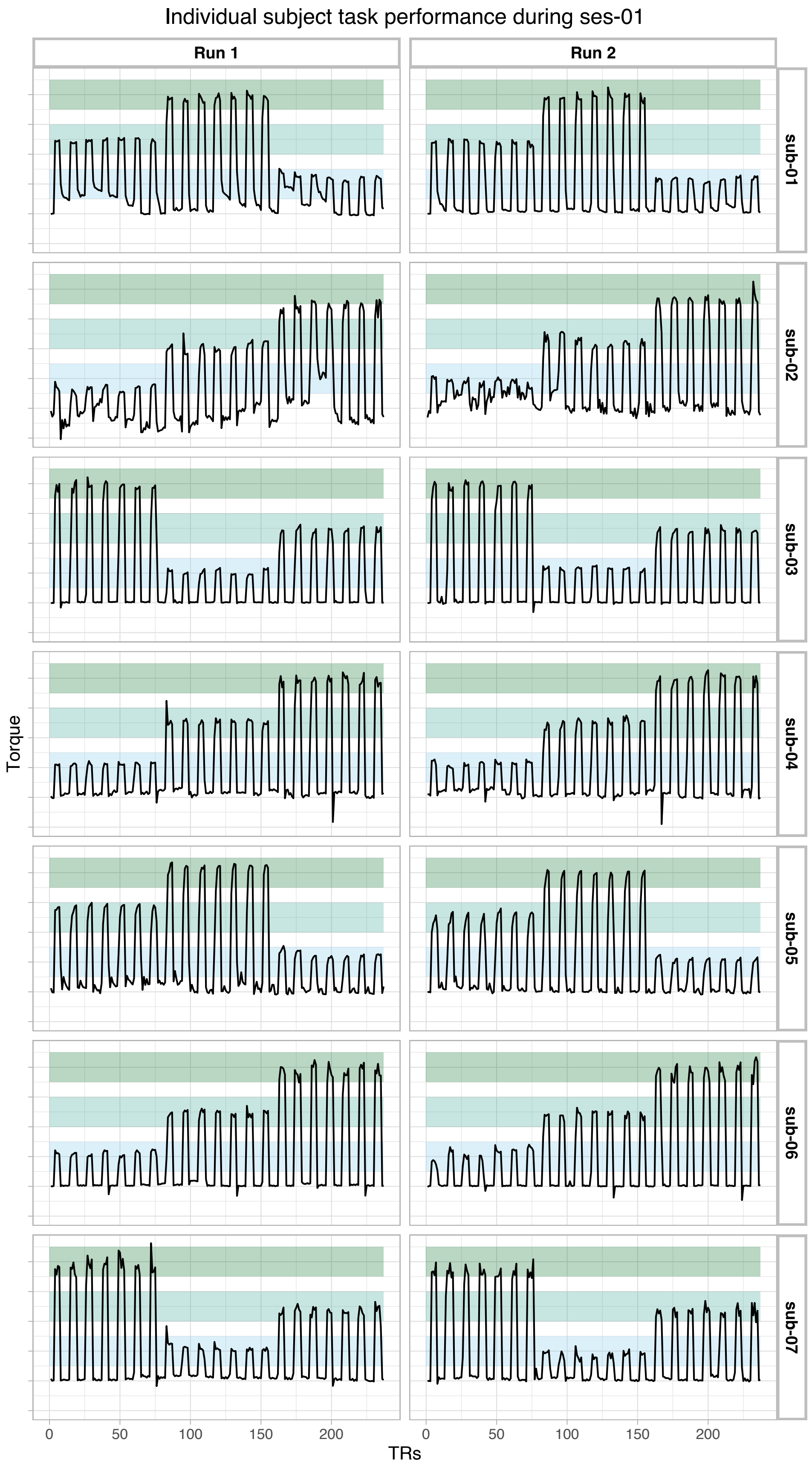
Individual subject task performance during ses-01
Run 1
Run 2
sub-01
sub-02
sub-03
sub-04
sub-05
sub-06
sub-07
Torque
TRs
0
50
100
150
200

Supplemental Figure 1. Individual subject task performance during session one, shown for runs 1 and 2. Torque targets were displayed to participants on a screen as part of real-time visual feedback on task performance. These torque targets are displayed in the figure as shaded blue/green: ± 5% target torque.

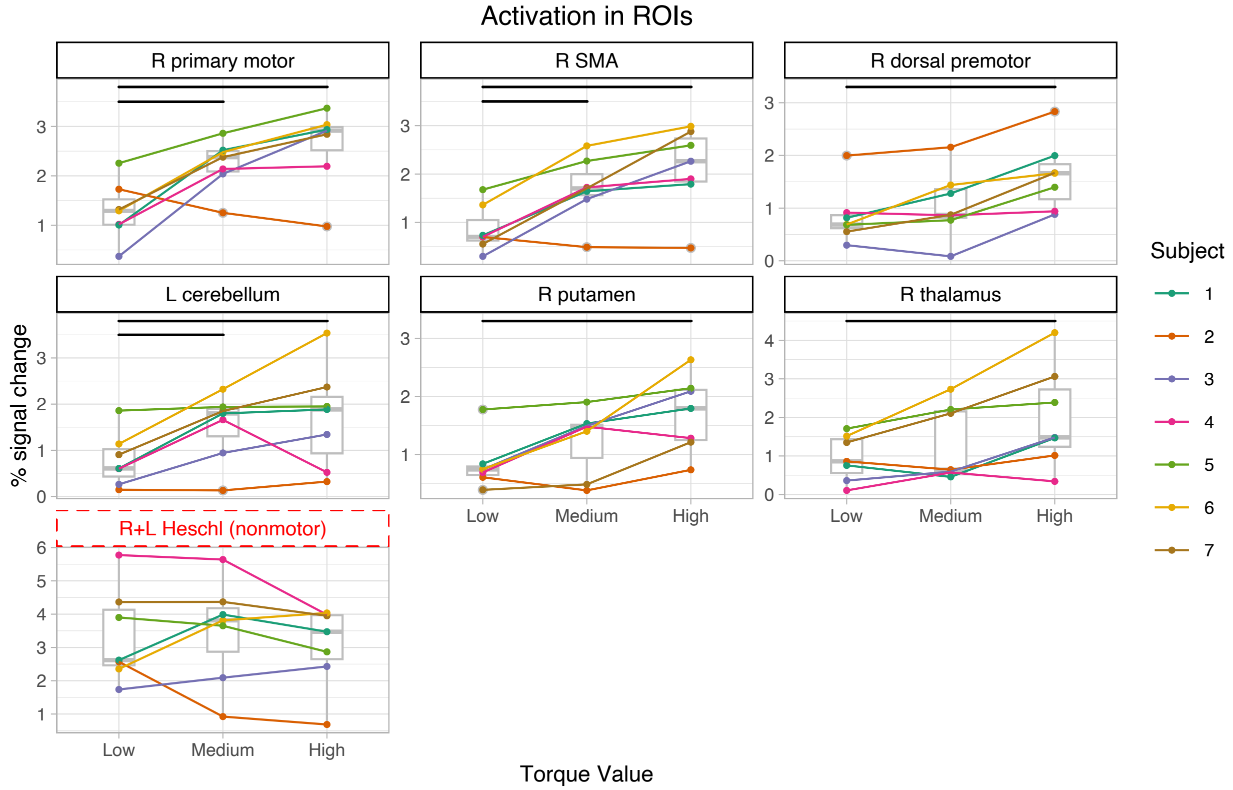


Supplemental Figure 2. Activation across torque levels in motor and non-motor ROIs. The top 10% of F-statistics in each motor region was calculated to create subject-specific ROIs. All motor ROIs had a significant effect of SABD (% MVT) on signal change (BetaCoefficient ~ SABD + (1|Subject)), with significant differences between Torque levels shown with a horizontal black bar. There was no significant effect of SABD on signal change in the Heschl's gyrus, related to auditory activation, which was expected to have consistent activation across all motor tasks.

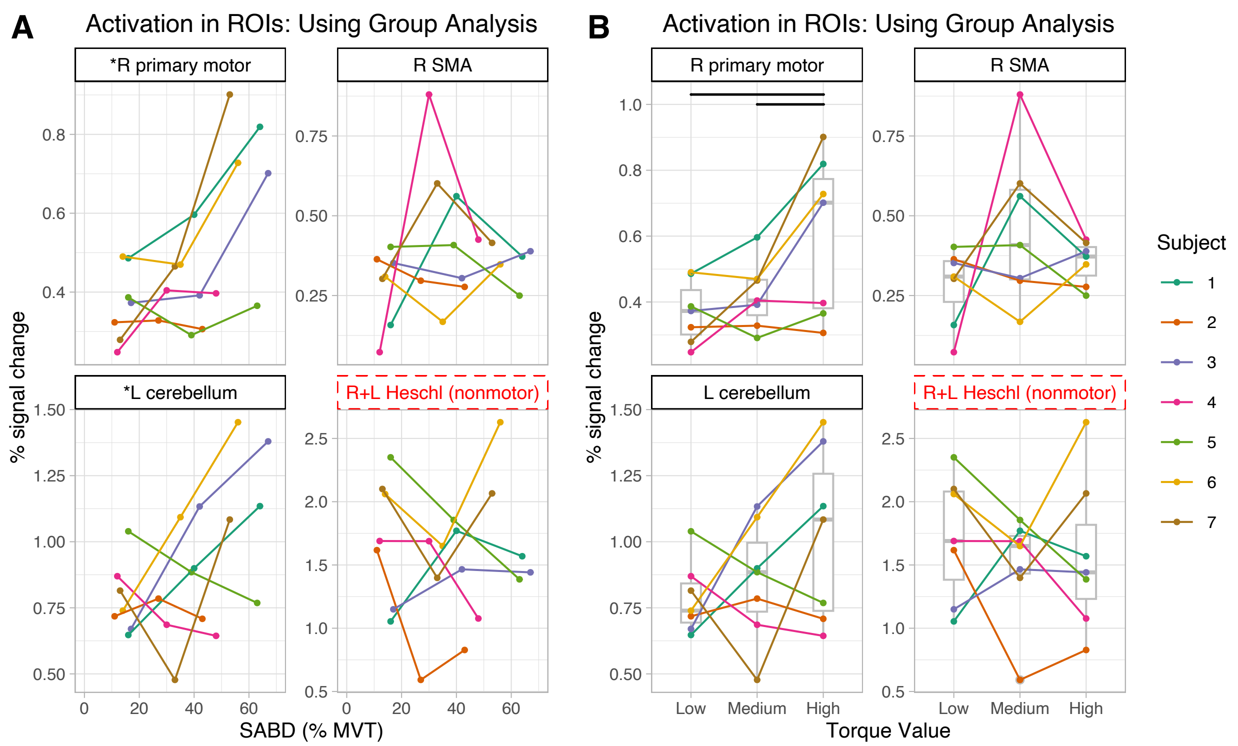


Supplemental Figure 3. Activation across torque levels in motor and non-motor ROIs, with ROIs created from group-level results. Activation is shown plotted against (A) shoulder abduction in % MVT and (B) Low, Medium, and High torque levels. The right primary motor cortex and left cerebellum had a significant effect of SABD (% MVT) on signal change, indicated by an asterisk next to the ROI names (BetaCoefficient ~ SABD + (1|Subject)). Significant differences between Torque levels are shown with a horizontal black bar in (B).

### Group analysis with each torque as a separate input

The incorporation of only seven subjects in our group analysis may limit power and reduce resulting group-level clusters. Therefore, we tested an identical group analysis to that described in the Methods, now incorporating data from each torque as a separate input to increase the apparent sample size (3 inputs per subject, 21 total inputs to group analysis). This method resulted in more group-level clusters, including clusters in the PMd and putamen (Supplemental Figure 4). There were still no thalamus clusters detected with this method.

As described with our main group analysis, we similarly created ROIs from group-level clusters in each region and used these ROIs to calculate shoulder motor activation for each subject and torque level (Supplemental Figure 5). We found a significant effect of shoulder abduction on torque level in M1 ($p < 0.005$), PMd ($p < 0.05$), cerebellum ($p < 0.05$), and putamen ($p < 0.005$). We did not find a significant effect of shoulder abduction on torque level in SMA ($p = 0.065$) or Heschl's gyrus ($p = 0.5798$).

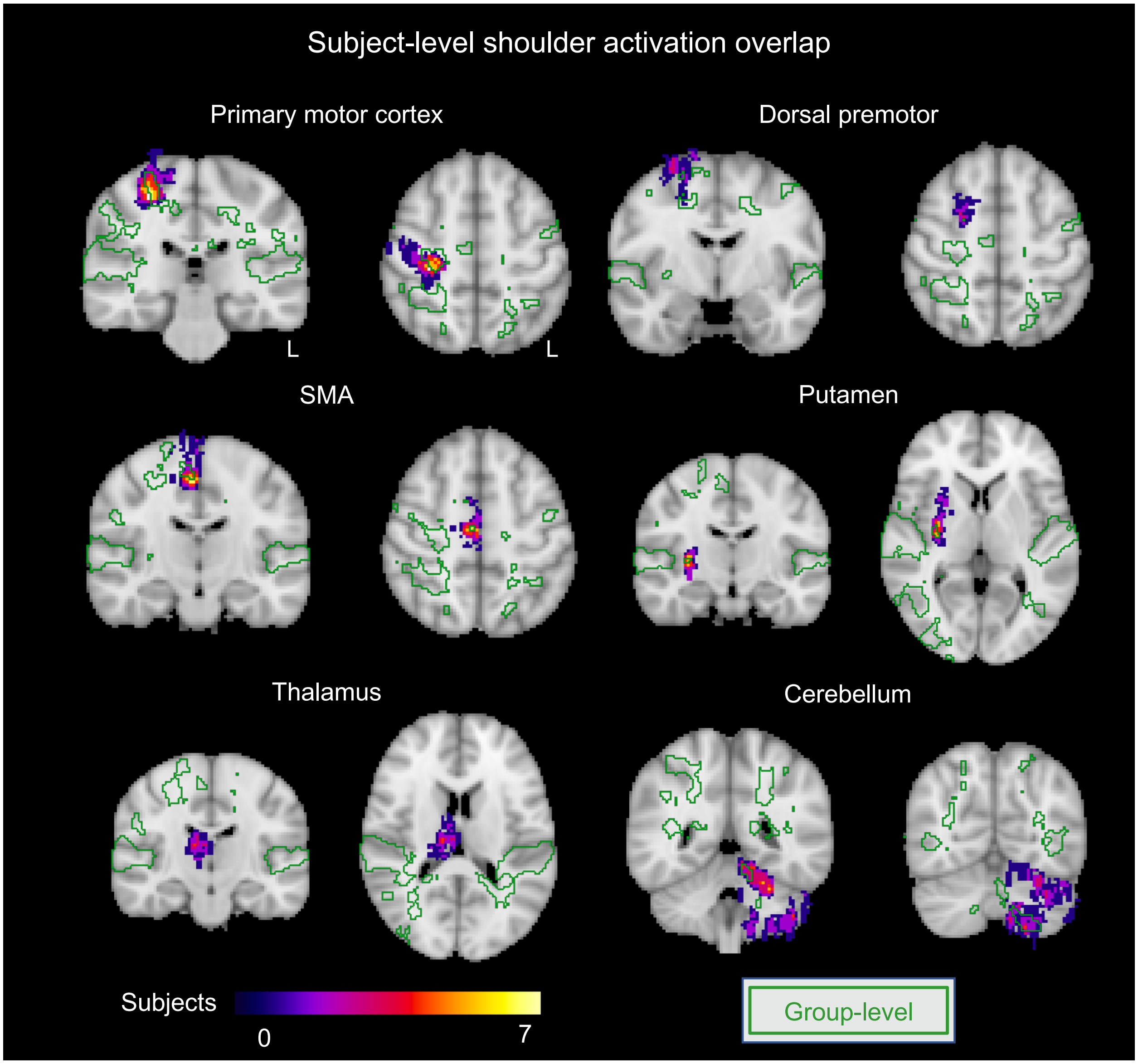


Supplemental Figure 4. Subject-level activation related to shoulder motor activity. Seven subjects were included. The top 10% of F-statistics in each ROI were transformed to MNI space for each subject, then results from all subjects were overlaid. For comparison, the group-level results for shoulder motor activity are shown in green outline; the visualized group-level results incorporated inputs from each torque level separately (21 total inputs).

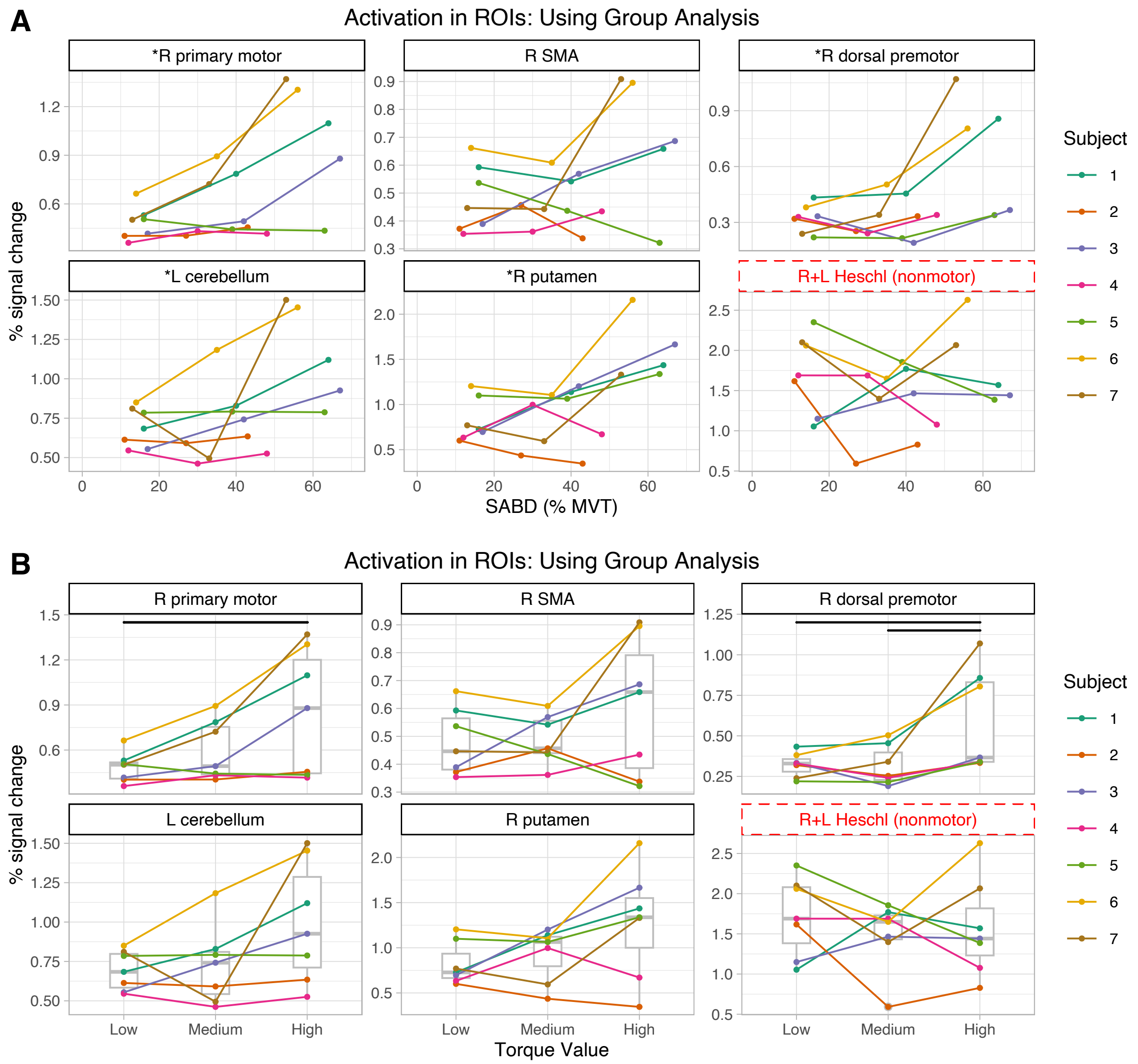


Supplemental Figure 5. Activation across torque levels in motor and non-motor ROIs, with ROIs created from group-level results that incorporated inputs from each torque level separately (21 total inputs). Activation is shown plotted against (A) shoulder abduction in % MVT and (B) Low, Medium, and High torque levels. The right primary motor cortex, right dorsal premotor region, right putamen, and left cerebellum had a significant effect of SABD (% MVT) on signal change, indicated by an asterisk next to the ROI name (BetaCoefficient ~ SABD + (1|Subject)). Significant differences between Torque levels are shown with a horizontal black bar in (B).

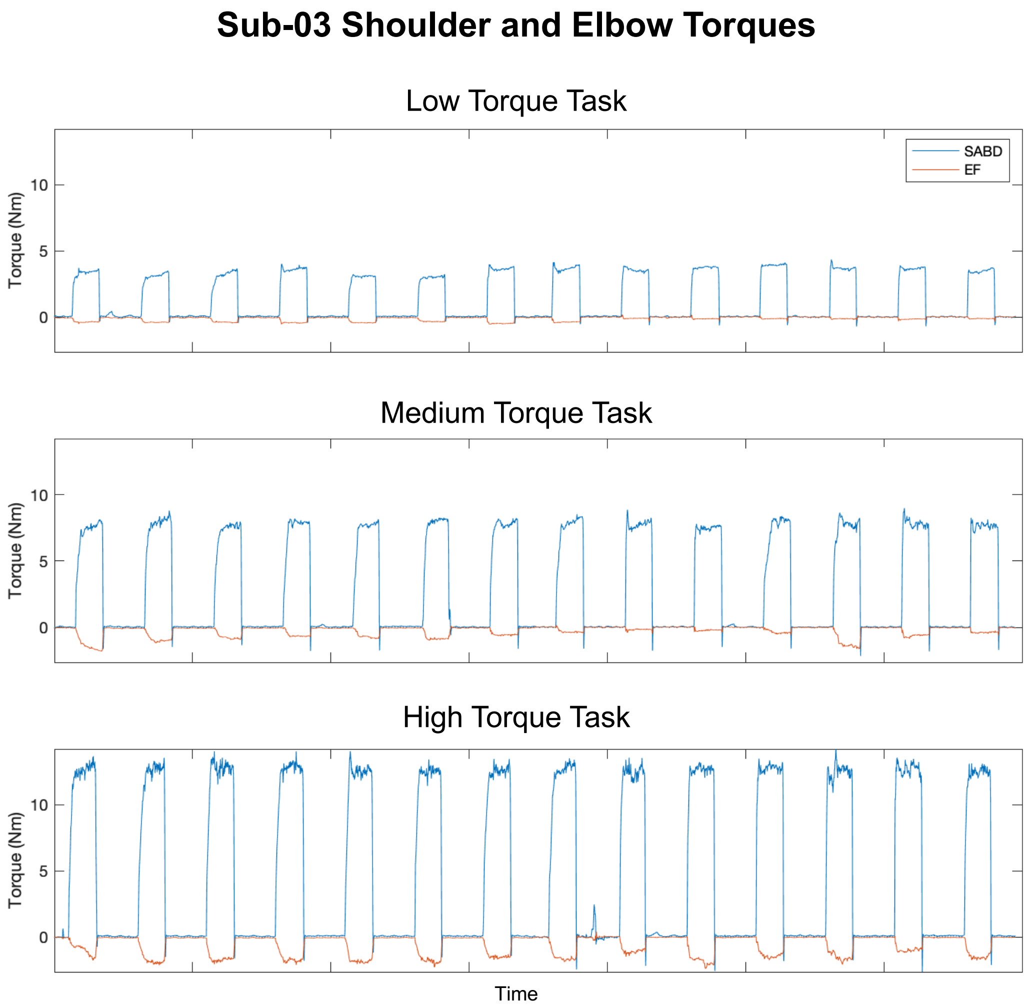


Supplemental Figure 6. Example shoulder abduction (SABD) and elbow flexion (EF) torques recorded from sub-03's performance of the Low, Medium, and High torque tasks in MRI session one. This subject demonstrated concurrent elbow extension during periods of shoulder abduction, with the magnitude of elbow extension increasing with magnitude of shoulder abduction.

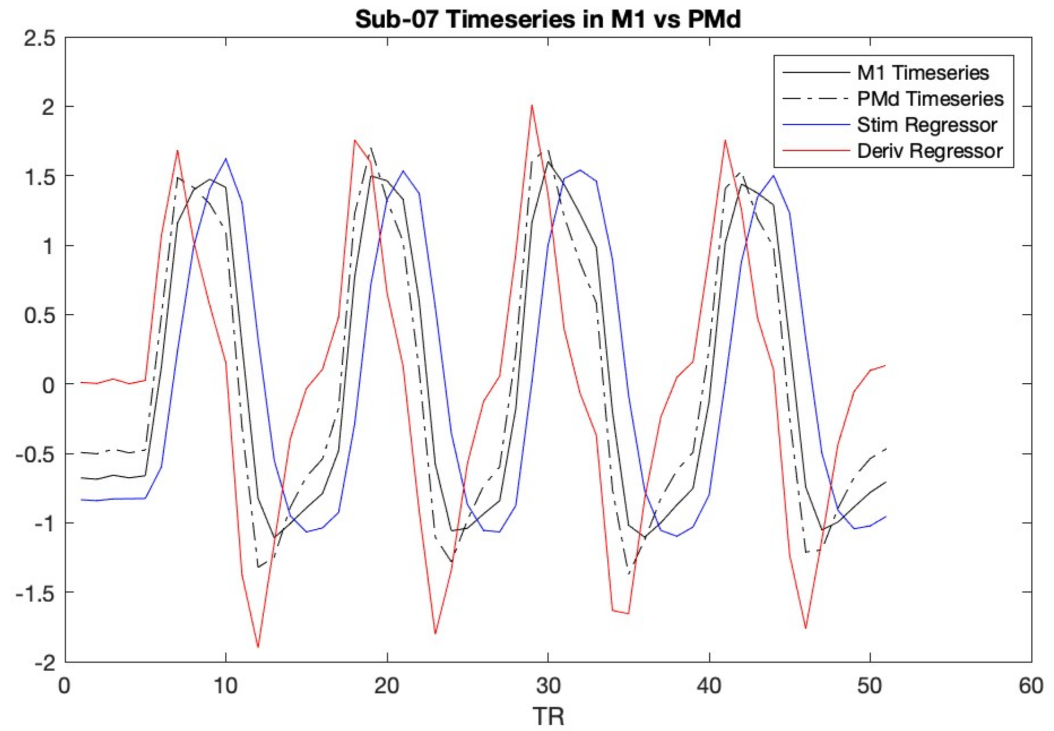


Supplemental Figure 7. Example of motor response variability in one subject. The mean combined motor timeseries within the subject-specific M1 and PMd ROIs were calculated and plotted against the motor-task stimulus regressor and its derivative. The PMd timeseries is slightly earlier than the M1 timeseries and more closely resembles the derivative compared to the motor-task stimulus.
